# Cholinergic neuromodulation of inhibitory interneurons facilitates functional integration in whole-brain models

**DOI:** 10.1101/2020.09.04.282095

**Authors:** Carlos Coronel-Oliveros, Rodrigo Cofré, Patricio Orio

## Abstract

Segregation and integration are two fundamental principles of brain structural and functional organization. Neuroimaging studies have shown that the brain transits between different functionally segregated and integrated states, and neuromodulatory systems have been proposed as key to facilitate these transitions. Although computational models have reproduced the effect of neuromodulation at the whole-brain level, the role of local inhibitory circuits and their cholinergic modulation has not been studied. In this article, we consider a Jansen & Rit whole-brain model in a network interconnected using a human connectome, and study the influence of the cholinergic and noradrenergic neuromodulatory systems on the segregation/integration balance. In our model, a newly introduced local inhibitory feedback enables the integration of whole-brain activity, and its modulation interacts with the other neuromodulatory influences to facilitate the transit between different functional states. Moreover, the new proposed model is able to reproduce an inverted-U relationship between noradrenergic modulation and network integration. Our work proposes a new possible mechanism behind segregation and integration in the brain.

## Introduction

Integration and Segregation of brain activity are nowadays two well-established brain organization principles [1–4]. Functional segregation refers to the existence of specialized brain regions, allowing the local processing of information. Integration coordinates these local activities in order to produce a coherent response to complex tasks or environmental contexts [1,2]. Both segregation and integration are required for the coherent global functioning of the brain; the balance between them constitutes a key element for cognitive flexibility, as highlighted by the theory of coordination dynamics [5,6].

From a structural point of view, the complex functional organization of the brain is possible thanks to an anatomical connectivity that combines both integrated and segregated network characteristics, having small-world and modular properties [7]. In spite of this *structural connectivity* (SC) remaining fixed over short time scales, different patterns of *functional connectivity* (FC) can be observed during the execution of particular behavioral tasks [2]. Moreover, functional Magnetic Resonance Imaging (fMRI) neuroimaging studies show that during a resting state the FC is not static, but rather evolves over the recording time. The non-stationarity of functional connectivity, referred as Functional Connectivity Dynamics (FCD), captures the variable nature of the brain dynamics [8,9]. In this context, an interesting question emerges: *How does the brain manage to produce dynamical transitions between different functional states from a rigid anatomical backbone?.*

Neuromodulatory systems provide a biophysical mechanism that enhances the dynamical flexibility. It is thought that functional integration arises by the increase of excitability and selectivity of neuronal populations, and a recent hypothesis proposed by [10] argues that neuromodulation allows the transition between integrated and segregated states, manipulating the neural gain function [11]. Indeed, the cholinergic system increments the overall excitability [12,13], and consequently rises population activity above noise, a mechanism referred as response gain [11]. The increase in signal-to-noise ratio, especially in brain areas that are close to each other, promotes segregation when considering the response gain by itself [10]. On the other hand, the noradrenergic system increases the reponsivity (or selectivity) of neuronal populations to input-driven activity respect to spontaneous activity [14–16] and filters out noise [17], a mechanism called filter gain [11]. This effect is more pronounced between distant brain regions, in which structural connectivity is relatively low, promoting functional integration [10]. In reality, a complex interaction between the cholinergic and noradrenergic system seems to manage the balance between integration and segregation. Using a whole-brain model, [18] showed that the inverted-U relationship between neuromodulation and integration, which has been reported in the literature [19], can be reproduced manipulating the effects of cholinergic and noradrenergic systems on neural gain. In this model, integration is not promoted by any neuromodulatory system by itself, but emerges by the combined action of both systems: the cholinergic and noradrenergic systems do not operate in an antagonistic fashion.

There are still unanswered questions about the specific effects of neuromodulation on integration and segregation. The overall increase of excitability, mediated by the cholinergic system, affects both the excitatory and inhibitory neuronal populations, effect well known at the meso-scale level [11,20–23]. Experimental research points out that the cholinergic system, through both nicotinic and muscarinic receptors, boosts the signal-to-noise ratio in two principal ways [11,22]: first, increasing the excitability of pyramidal neurons [23–25], and second, enhancing the activity and firing rates of dendritic-targeting GABAergic interneurons, an effect that promotes intra-columnar inhibition, reducing the local excitatory feedback to pyramidal neurons [23, 26, 27]. Consequently, pyramidal neurons become more responsive to stimulus from other distant regions respect to the stimulus of its own cortical column [21,22,24]. The particular effect of the cholinergic system in excitatory neurons was one of the focus of the whole-brain simulation work by [18]. However, the cholinergic modulation of inhibitory interneurons and its effect on the segregation/integration balance has not been analyzed at the whole-brain level, and comprises the main focus of the present work.

Here, we use an *in silico* approach to analyze the effect of neuromodulatory systems on functional integration in the brain, focusing on the cholinergic action in inhibitory interneurons. Whole-brain computational models can reproduce statistical features of neuroimaging signals from human brains [28,29], providing a tool to explore the computational and biophysical mechanisms that underlie the organization principles of integration and segregation [18,30,31]. We combined a real human structural connectivity with the Jansen & Rit neural mass model of cortical columns [32], widely used to reproduce electroencephalography (EEG) signals in the healthy and pathological brain [33–35]. fMRI-blood-oxygen-level dependent (BOLD) signals were generated from the firing rates of pyramidal neurons to quantify integration and segregation in the functional connectivity matrices derived from the BOLD-like signals using a graph theoretical approach.

The neuromodulation was discerned in three components. First, we included an “excitatory gain”, which increases the inter-columnar coupling. This gain mechanism is mediated by the action of the cholinergic system in pyramidal neurons, principally but not exclusively, and increments pyramidal excitability [10,11,22]. Second, we added an “inhibitory gain”, also mediated by the cholinergic system, that controls the inputs from inhibitory to excitatory interneurons and reduces the local feedback excitation. This additional connection, well described in cortical columns [36, 37], represents a modification of the original neural mass model proposed by [32]. Finally, we incorporated a “filter gain”, that increments the pyramidal neurons sigmoid function slope [11]. This last gain mechanism is mediated by the noradrenergic system; it acts as a filter, decreasing (increasing) the responsivity to weak (strong) stimuli [15,17], boosting signal-to-noise ratio and promoting integration [10].

Our results show, in the context of a whole-brain model, that the control of local feedback excitation, mediated by the action of the cholinergic system on inhibitory interneurons, is necessary for the modulation of the segregation/integration balance by the other systems. This constitutes a step forward from the neuromodulatory framework proposed by [10], including the role of a second cholinergic target and also highlighting the role of a homeostatic inhibitory feedback. We also describe that integration is accompanied by an increment in the signal-to-noise ratio, and a reduction of dynamical variability captured by the FCD analysis. Our work sheds light in how meso-scale properties, such as local inhibition, shape statistical network features at the macro-scale level.

## Results

We assessed the effect of the neuromodulatory systems using a whole-brain neural mass model of brain activity. In the model, each node corresponds to a brain area and is represented by a neural mass consisting of three populations [32]: pyramidal neurons, excitatory interneurons, and inhibitory interneurons (Fig 1A). Based on Silberberg & Markram [36] and Fino *et al.* [37], we have added a connection from inhibitory interneurons from excitatory interneurons (dotted line in Fig 1A), allowing us to study the effect of its modulation by cholinergic influences (see below). The nodes are connected through a weighted and undirected structural connectivity matrix derived from human data [38], parcellated in 90 cortical and sub-cortical regions with the automated anatomical labeling (AAL) atlas [39]. Further, we included heterogeneous time delays based in the spatial location of brain regions defined by AAL parcellation [39] (Fig 1B). Connections between nodes are made by pyramidal neurons, considering that long-range projections are mainly excitatory [40,41]. Using the firing rates of each node as inputs to a generalized hemodynamic model [42], we obtained fMRI-BOLD signals from which we calculated integration and segregation of the resulting functional connectivity matrices.

**Fig 1.**
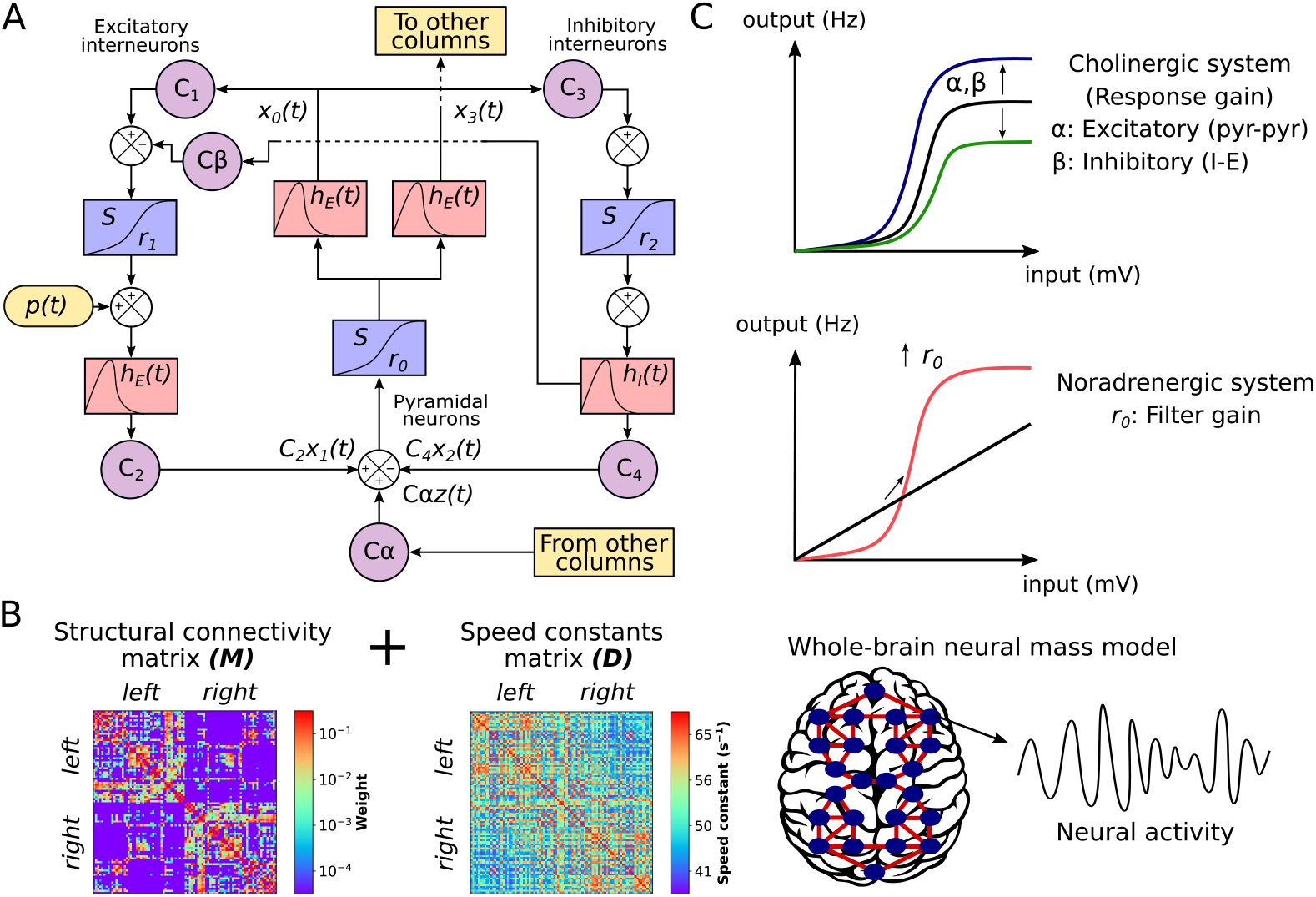
Whole-brain neural mass model. **A)** The Jansen & Rit model is constituted by a population of pyramidal neurons with excitatory and inhibitory feedback mediated by interneurons (INs). Each population is connected by a series of constants *C_i_*. The outputs are transformed from average pulse density to average postsynaptic membrane potential by an excitatory (inhibitory) impulse response function *h_E_*(*t*) (*h_I_*(*t*)). Then, a sigmoid function *S* performs the inverse operation. Pyramidal neurons project to distant cortical columns, and receive both uncorrelated Gaussian-distributed inputs *p*(*t*) and inputs from other cortical columns *z*(*t*). Neuromodulation is constituted by the excitatory gain *α*, which scales *z*(*t*), inhibitory gain *β*, which increases the inhibitory input to excitatory INs, and filter gain, *r*_0_, which modifies the slope of the sigmoid function in pyramidal neurons. **B)** In the whole-brain model, each node represents a cortical column, whose dynamics is ruled by the Jansen & Rit equations. Nodes are connected through a structural connectivity matrix, *M*, and a speed constants matrix, *D.* **C)** Neuromodulation modifies the input (average postsynaptic membrane potential) to output (average pulse density) sigmoid function. The cholinergic system has a multiplicative effect on the sigmoid function. *α* amplifies the response of pyramidal neurons to other columns’ input, while *β* amplifies the effect of inhibitory INs to excitatory INs. On the other hand, the noradrenergic system increments the responsivity of pyramidal neurons to relevant stimuli respect to noise, as a filter, by increasing the slope *r*_0_ of their sigmoid function.

Following Shine *et al.* [18], we modeled the influence of the cholinergic and noradrenergic systems through the manipulation of the response and filter gain, respectively (Fig 1C). The principal difference in our approach is that we split the response gain in excitatory gain (long-range pyramidal to pyramidal coupling), *α*, and inhibitory gain (local inhibitory to excitatory interneurons coupling), *β.* While the excitatory gain boosts pyramidal neurons output, the inhibitory gain reduces the local excitatory feedback from interneurons. Finally, the filter gain *r*_0_ modifies the sigmoid function slope of pyramidal neurons, increasing its responsivity to relevant stimuli and boosting signal-to-noise ratio. Here, we studied the combined effect of the three gain mechanisms to understand how neuromodulatory systems shape the global neuronal dynamics in two different timescales: EEG-like and BOLD-like signals. Our hypothesis is that the inhibitory gain will play a significant role in increasing the likelihood of integration.

### Inhibitory gain facilitates neuronal coordination

We first studied the combined influence of the excitatory and inhibitory response gains, by fixing *r*_0_ = 0.56 mV^−1^ and then simulating neuronal activity at different combinations of *α* ∈ [0,1] and *β* ∈ [0,0.5]. Then, we measured the degree of synchrony of the EEG-like signals using the averaged Kuramoto order parameter 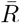 [43]. In addition, we analyzed the graph properties of the *static* (time-averaged) functional connectivity (sFC) matrices, obtained from the pairwise Pearson’s correlations of BOLD-like signals. Namely, we calculated the global efficiency *E^w^*, a measure of integration defined as the inverse of the characteristic path length [44], and modularity *Q^w^*, a measure of segregation based on the detection of network communities or modules [44]. High values of *E^w^* represent an efficient coordination between all pairs of nodes in the network, a signature of integration. In contrast, a high modularity *Q^w^* is associated to segregation and viceversa [44]. Finally, we measured the mean participation coefficient *PC*^w^, an integration metric that quantifies the between-modules connectivity, and the transitivity *T*^w^, which accounts for segregation counting triangular motifs [44].

Fig 2A shows that functional integration (*E^w^*) and neuronal synchronization 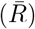 are maximized in an intermediate region of the (*α, β*) parameter space. Also, coordination is accompanied by a decrease in the mean oscillatory frequency, *ω*, which falls within the EEG-Theta range (4-8 Hz), and a decrease in the segregation (*Q^w^*). The system undergoes a sharp transition crossing a critical boundary both in EEG and BOLD timescales. This clear delimitation of states is a feature of criticality [45]. The transitions between different regimes are better appreciated in Fig 2B, where we show a 1-D sweep of *α* at *β* = 0.25. Dashed lines at *α* = 0.23 and *α* = 0.8 correspond to points in the parameter space where drastic changes in dynamic properties of the network occur. Both *E^w^* and 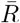 follow an inverted-U relationship with excitatory neuromodulation, recovering a known result in this field [19]. Also, *Q^w^* peaks higher at the right critical boundary (dashed lines), supporting the hypothesis that further increases of the excitatory gain, mediated by the cholinergic system, promote segregation [10]. A 1-D sweep of *β* at *α* = 0.5 (Fig 2C), shows an increase in synchronization and integration crossing the critical point at *β* = 0.1. The aforementioned results are similar for the mean *PC^w^* and *T^w^*, as shown in S1 Fig.

**Fig 2.**
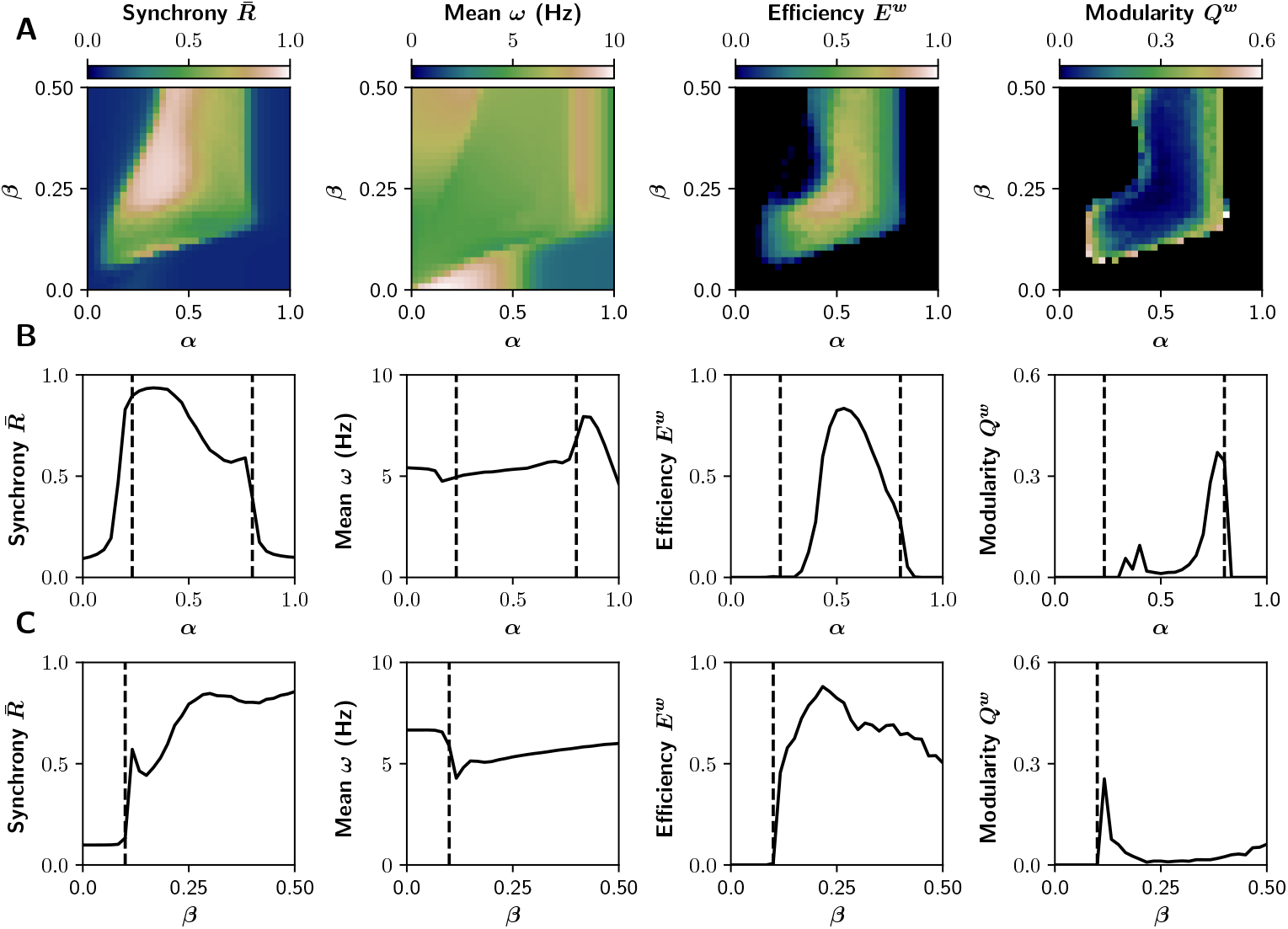
Signal and network features in the (*α,β*) parameter space. **A)** Average phase synchrony 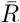, mean oscillatory frequency *ω* of EEG-like signals, global efficiency *E^w^* (integration) and modularity *Q^w^* (segregation) of the graphs derived from the sFCs of the BOLD-like signals. **B)** Transitions through critical boundaries in the direction of *α* axis, for a fixed *β* = 0.25. Transition points are represented by black dashed lines at *α* = 0.23 and *α* = 0.8. **C)** Transitions in the direction of *β* axis, for a fixed *α* = 0.5, with a critical point at *β* = 0.1.

The modulation of the inhibitory gain (*β*) shows a compelling effect on the integration and segregation of the whole network. This could be due to the reduction of excitatory feedback only, or a more specific effect of the connection from inhibitory to excitatory interneurons. In the first case, we expect a similar effect by reducing the *C*_1_ parameter (see Fig 1A) because this also reduces the excitatory feedback loop of the cortical columns. As shown in S2 Fig, this is only partially the case. The reduction of the *C*_1_ connection weight –in the absence of the inhibitory-to-excitatory interneuron connection– enables the network to reach integration but in a smaller region of the parameter space and to a lower extent than the inhibitory modulation that we introduced in our model. This highlights the role of specific intra-columnar inhibitory feedback connections in shaping the network behavior, and justifies our modification of the model as an homeostatic mechanism (see Discussion).

The reduction of the average oscillatory frequency could be an effect of the time delays incorporated in the model, as suggested by Nordenfelt *et al.* [46] and Lea-Carnall *et al.* [47]. We found the mean *ω* to be a function of the average speed constant 〈*D*〉 (not shown), suggesting that whole-brain integration goes together with a reduction in oscillatory frequency, as a consequence of the time delays between cortical columns.

At the functional level, integration is characterized by the increase of inter-modular connectivity. To observe in detail how each gain mechanism produces integrated or segregated states, we show some BOLD-like signals and their respective sFCs matrices in Fig 3. We chose five tuples of values of (*α,β*), marked with the red circles in the Fig 3A. We observe that functional integration is maximal in the middle (*α* = 0.5, *β* = 0.25), and segregation is promoted far away from this point (*α* = 0.25, *β* = 0.125, and *α* = 0.75, *β* = 0.375). In the extreme cases (*α* = 0, *β* = 0, and *α* = 1, *β* = 0.5) there is neither integration nor segregation.

**Fig 3.**
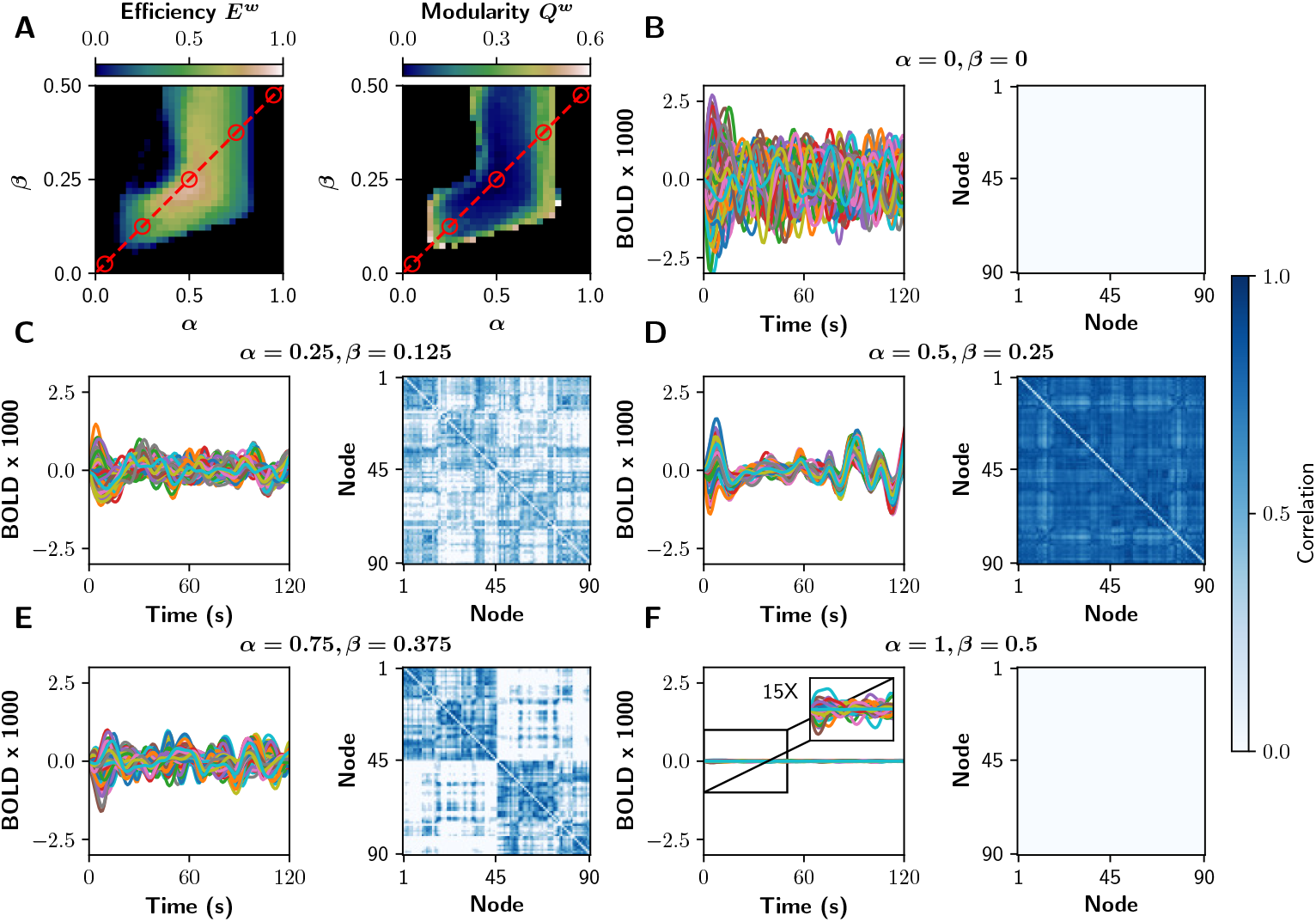
fMRI-like sFCs at different values of *α* and *β*. **A)** The red circles represent pairs of (*α,β*) values in which different integration/segregation profiles can be observed. **B-F)** BOLD-like signals, and their respective sFCs matrices, for the (*α, β*) shown in A. We shown the first 120 s of BOLD-like signals, while sFCs matrices were built with the full-length time series (600 s).

### Inhibitory gain allows the noradrenaline-mediated integration

The inclusion of the inhibitory gain in the model has the capability of producing novel predictions on the basis of a biophysically plausible mechanism. To validate this model and its results, it should also reproduce the results of the current neuromodulatory paradigm proposed by Shine [10,18]. We characterized the relationship between neuromodulation and integration in the (*α, r*_0_) parameter space, with *α* ∈ [0,1] and *r*_0_ ∈ [0,1] while leaving *β* fixed at 0 or 0.4 (without and with inhibitory gain, respectively). The results for *β* = 0 (Fig 4A) show neither integration nor synchronization in the entire parameter space. On the other hand, the observations of Shine *et al.* [18] are fully reproduced with *β* = 0.4 (Fig 4B), supporting the fact that inhibitory gain is essential to produce neuronal coordination. Similar results hold for the mean *PC^w^* and *T^w^*, as shown in S3 Fig.

**Fig 4.**
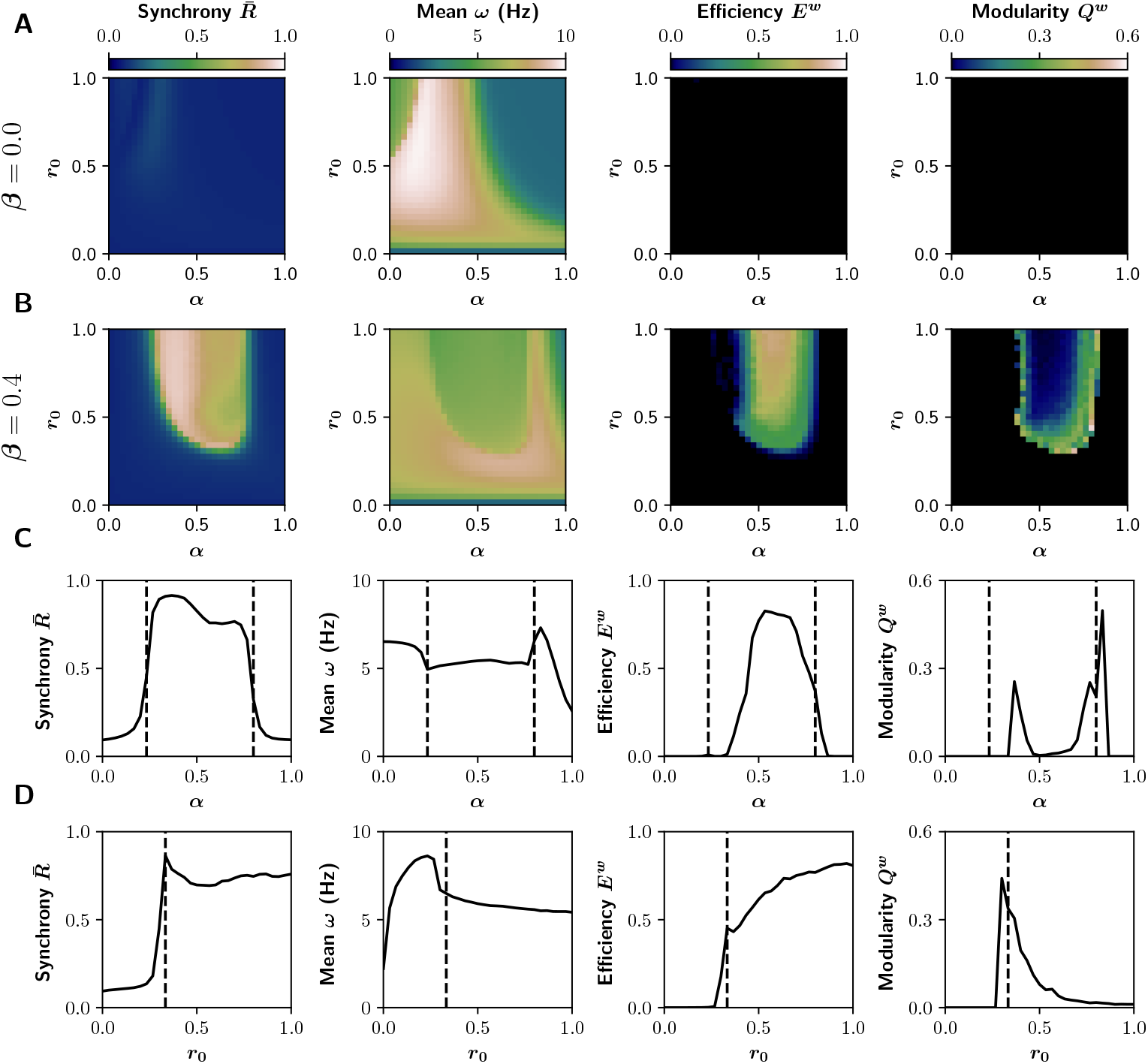
Signal and network features in the (*α, r*_0_) parameter space. **A-B)** Average phase synchrony R and mean oscillatory frequency *ω*, and global efficiency *E^w^* (integration) and modularity *Q^w^* (segregation) of the graphs derived from the sFCs of the BOLD-like signals, for **A)** *β* = 0 (no action of the inhibitory gain) and **B)** *β* = 0.4. **C)** Transitions through the critical boundary in the direction of *α* axis, with a fixed *r*_0_ = 1 mV^−1^ and *β* = 0.4. Critical transition points represented by black dashed lines at *α* = 0.23 and *α* = 0.8. **D)** Transitions in the direction of *r*_0_ axis, for a fixed *α* = 0.6, with a critical point at *r*_0_ = 0.33 mV^−1^ and *β* = 0.4.

As observed previously in Fig 2, a critical boundary delimits asynchronous and synchronous states in the (*α,r*_0_) parameter space, a signature of criticality [45]. This behavior was reported before in the original neuromodulatory framework of Shine *et al.* [18]. A 1-D sweep of *α* at *r*_0_ = 1 mV^−1^ shows a sharp transition (Fig 4C). Mean phase synchrony 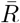 increments alongside global efficiency *E^w^*, changes that go along with a decrease of the average frequency *ω*, and a reduction of modularity *Q^w^*. However, further increments of *α* produce network desynchronization. On the other hand, a 1-D sweep of *r*_0_ at *α* = 0.6 (Fig 4D) produces similar observations, but just one boundary is visible. These results support the hypothesis that noradrenergic and cholinergic systems, combined, promote integration following an inverted-U relationship [4,11,18], but only if the inhibitory gain is included.

In the whole-brain model, the cholinergic system exerts its effect by changing both *α* and *β* parameters. Under this assumption, a logical consequence of the cholinergic neuromodulation is the possibility of simultaneously increase/decrease of *α* and *β*. For that reason, we repeated the analysis previously performed in the (*α*, *r*_0_) parameter space, but this time we changed *β* alongside *α* following the relationship *β* = 0.35*α*. The results are shown in S4 Fig. They are similar to those in Fig 4, but this time the relationship between *r*_0_ and *E^w^* is no longer a sigmoid-like function, rather it follows an inverted-U relationship. The simultaneously modulation of *α* and *β* has a consequence to the excitability levels of pyramidal cells. For example, when *α* ~ 0.5 the maximum integration could be achieved at lowers levels of *r*_0_. Increasing *r*_0_ further produces an over-excitation of pyramidal cells. In consequence, for *α* = 0.5, the optimal *r*_0_ value for maximizing integration in S4 Fig (with *β* = 0.35*α* = 0.175) would be lower in comparison with Fig 4 (with a fixed *β* = 0.4). Thus, the simultaneous modulation of both excitatory and inhibitory circuits in a whole brain model, allowed to recover an empirical feature that was not previously reproduced [19].

### Signal-to-noise ratio and regularity matches with neuronal coordination

Previous experimental and theoretical works [10,11,22] suggest that neuromodulatory systems increase the signal-to-noise ratio, allowing neuronal populations to be sensitive to local or distant populations to a greater extent than noise. To test that, we measured the signal-to noise-ratio (SNR) using the power spectral density (PSD) function of each signal (see Methods) and report the average value over all nodes. Additionally, we computed the regularity index [48], as a measure of signal periodicity. This metric is defined as the second absolute peak of the autocorrelation function and is bounded between 0 and 1, with 0 for purely chaotic or noisy signals, and 1 for perfectly periodic signals. We report the average overall nodes.

Both SNR and regularity match the region of neuronal synchronization and integration (Fig 5), supporting the idea that neuromodulatory systems promote integration by increasing SNR. These results are not possible without the action of inhibitory gain (*β* = 0, Fig 5A). As mentioned before, in conditions of reduced excitatory feedback, the excitatory response gain increases the output of pyramidal neurons above noise and, consequently, boosts SNR. Then, the increase of filter gain raises the sensitivity of neuronal populations to inter-columnar stimulation.

**Fig 5.**
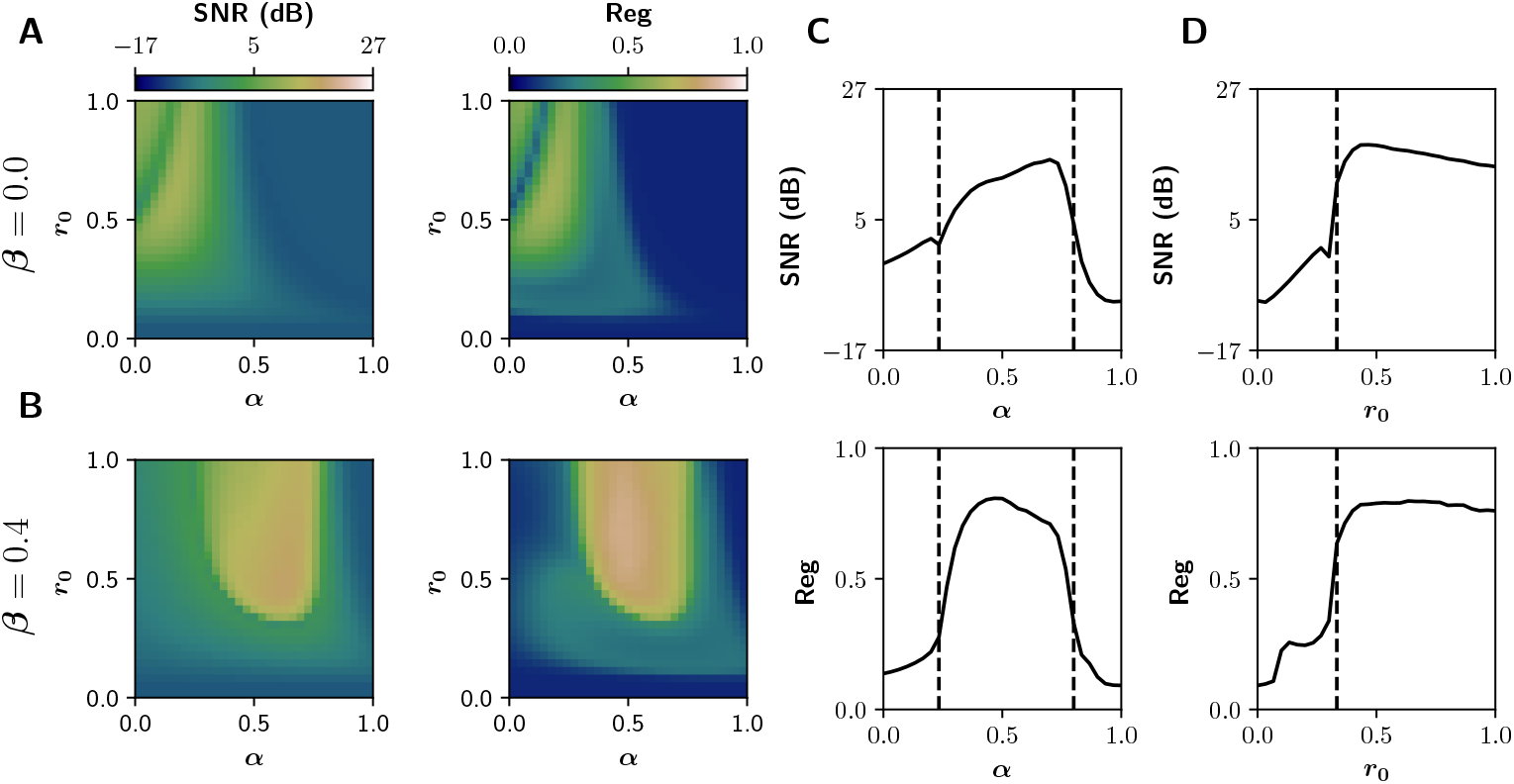
Signal-to-noise ratio (SNR) and regularity (Reg), for the EEG-like signals, in the (*α,r*_0_) parameter space. **A)** No action of inhibitory gain (*β* = 0). **B)** SNR and regularity matches with functional integration for *β* = 0.4. **C)** Transitions through the critical boundary in the direction of *α* axis, with a fixed *r*_0_ = 1 mV^−1^. Critical points represented by black dashed lines at *α* = 0.23 and *α* = 0.8. **D)** Transitions in the direction of *r*_0_ axis, for a fixed *α* = 0.6, with a critical point at *r*_0_ = 0.33 mV^−1^.

The increment in regularity suggests that a partial reduction of signal stochasticity is necessary for the whole-brain functional integration, a result supported by the idea that ordered in-phase synchronization is an essential and cost-efficient way to coordinate the brain activity [49]. Interestingly, regularity is near 0.5 at the critical boundary (Fig 5D), placing the signals between order and disorder, in which high amplitude synchronized oscillations alternate with noisy unsynchronized signals.

### Dynamical richness peaks near the critical boundary

We tested the hypothesis that network stability is higher in integrated states, and neuronal variability peaks near the critical boundaries. In the faster timescale (EEG), we computed the metastability *χ_R_* of bandpass filtered EEG-like signals, using a metric defined as the variance of the Kuramoto order parameter [50]. Values closer to 0 are expected for completely asynchronous or completely synchronous activity, and greater than 0 when the neuronal activity exhibits periods of high and low synchronization. In the slowest timescale, we performed a Functional Connectivity Dynamics (FCD) analysis [8,9] over the BOLD-like signals, using the sliding windows approach depicted in the Fig 6A-C [51]. The resulting time vs time FCD matrix captures the concurrence of FC patterns, visualized as square blocks. We computed the variance of the FCD, var(FCD), as a multistability index [51], where values greater than 0 indicate the switching between different FC patterns. Both metastability and multistability are measures of dynamical richness. Additionally, we calculated the FCD speed *d_typ_* as described by Battaglia *et al.* [52], which captures how fast the FC patterns fluctuate over time. Values closer to 1 indicate a recurrent change of diverse FC patterns, and closer to 0 the concurrence of stable and similar states over time.

**Fig 6.**
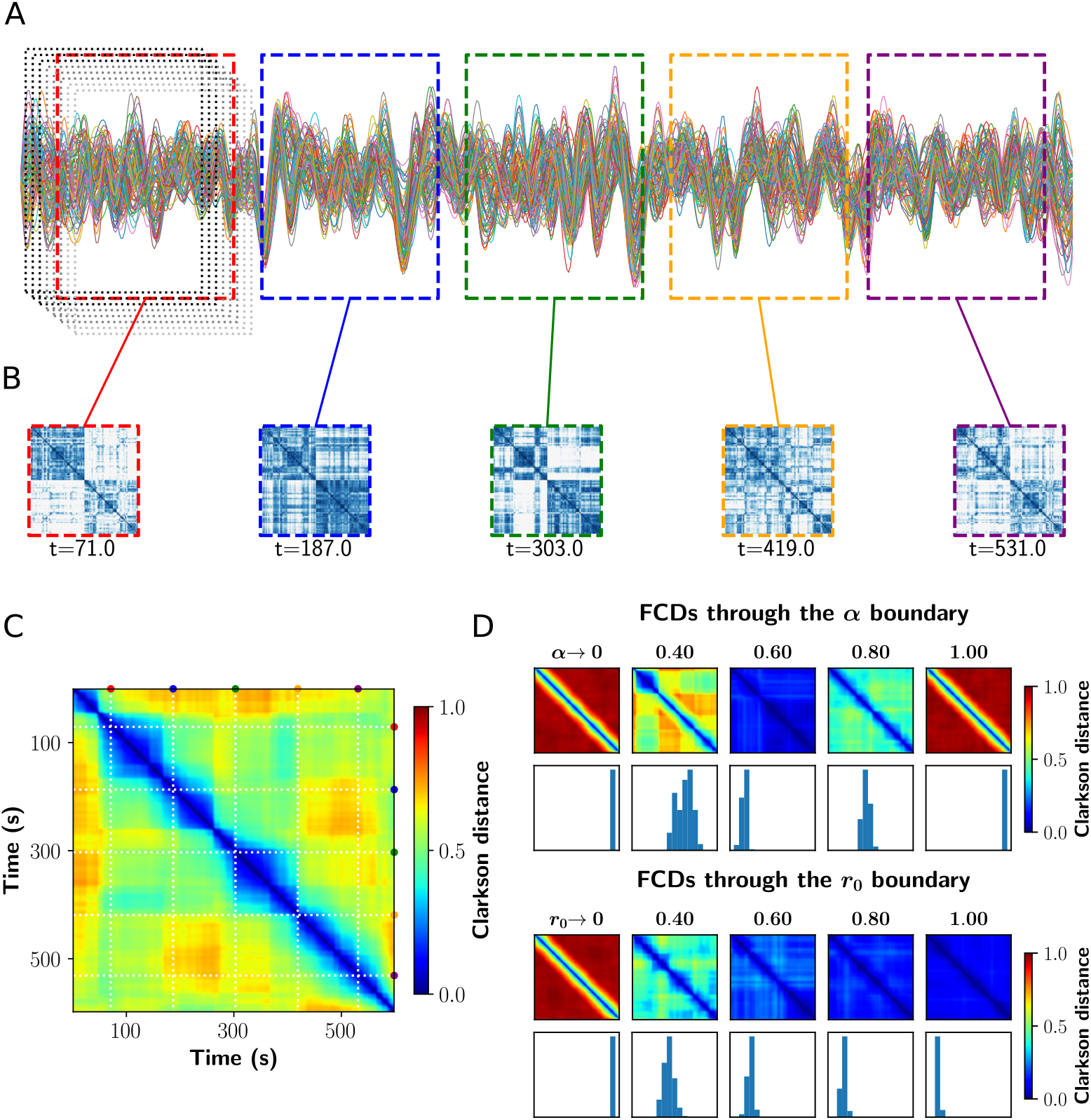
Analysis of Functional Connectivity Dynamics. **A)** Sample fMRI-BOLD time series showing the fixed length and overlapping time windows at the begginging. In color, the time windows corresponding to the FCs shown in B. **B)** FCs matrices obtained in the colored time windows. **C)** Functional Connectivity Dynamics (FCD) matrix, where all the FCs obtained were vectorized and then compared against each other using a vector-based distance (Clarkson distance). **D)** FCDs matrices through the critical boundary, in both *α* and *r*_0_ direction. Below each FCD, a histogram of its upper triangular values is shown. The variance of these values constitutes a measure of multistability.

In Fig 6D we show a set of FCD matrices obtained at different values of *α* and *r*_0_, together with histograms of their off-diagonal values. Red FCD matrices (with high values) correspond to incoherent states, as the FC continuously evolve in time. On the other hand, a blue FCD matrix (with low values) indicates a fixed FC throughout the simulation. Multistability is higher for green/yellow patchy matrices, because this indicates FC patterns that change and also repeat over time. As can be inferred observing the FCD distributions, the *variance* of the values in the histograms (var(FCD)) can be used as a measure of multistability [51].

Fig 7 shows how metastability, multistability (FCD variance) and FCD speed change in the whole (*α, r*_0_) space. At low levels of both *α* and *r*_0_, the neuronal activity is constituted mainly by noisy asynchronous signals, conditions associated to low (near 0) values of *χ_R_* and var(FCD), and with a high *d_typ_* (all FC patterns differ from each other, as expected for noise-driven signals) (Fig 7A). In the other extreme, for *r*_0_ > 0.5 mV^−1^ and *α* ∈ [0.4,0.6], values that correspond to the integrated states, var(FCD) is also small and *d_typ_* falls close to 0. In consequence, integrated states are more stable and less susceptible to network reconfiguration over time. In contrast, both *χ_R_* and var(FCD) peak near the critical boundary, through the direction of *α* and *r*_0_ axes (Fig 7B-C). Moreover, crossing the boundary is associated with a continuous decrease of *d_typ_*: the emerging integration mediated by gain mechanisms is associated with more stable FCs patterns over time. These observations are in agreement with experimental results using fMRI, in which the network variability increases in resting-state, and decreases with network integration during cognitive tasks [3].

**Fig 7.**
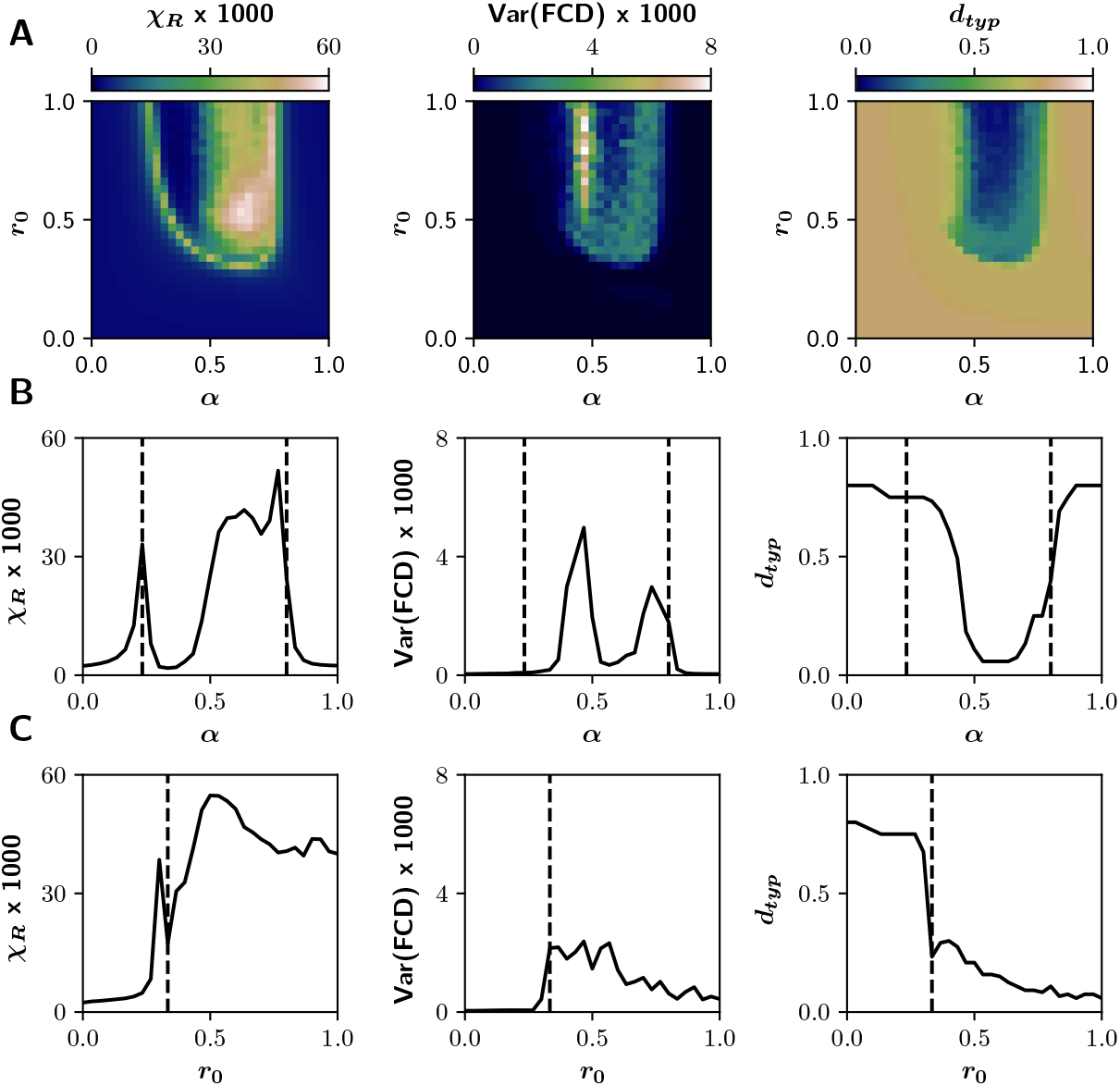
Dynamical features of the system in the (*α,r*_0_) parameter space. **A)** Metastability *χ_R_* calculated as the variance of the Kuramoto order parameter (EEG-like signals), multistability var(FCD) and typical FCD speed *d_typ_* measured from the Functional Connectivity Dynamics analysis (BOLD-like signals). While *χ_R_* and var(FCD) peak on the critical boundary, *d_typ_* decreases crossing the dashed lines (critical points) and reaches its minimum values in integrated states. **B)** Transitions through the critical boundary in the direction of *α* axis, with a fixed *r*_0_ = 1 mV^−1^. Critical points represented by black dashed lines at *α* = 0.23 and *α* = 0.8. **C)** Transitions in the direction of *r*_0_ axis, for a fixed *α* = 0.6, with a critical point at *r*_0_ = 0.33 mV^−1^.

## Discussion

Based on previous experimental findings [20, 23, 26, 27], we hypothesized that the cholinergic neuromodulation of the inhibitory interneurons (that suppresses the local excitatory feedback to pyramidal neurons) facilitates functional integration. Our results confirm this hypothesis in the context of a whole-brain neural mass model, and hold simultaneously for EEG-like and fMRI-like signals obtained considering different time scales. The main novelty of our work is the inclusion of the inhibitory gain (the parameter *β*) in the model and the characterization of its influence in the functional integration.

Several clues suggest that the model we proposed is in the right track. First, our results reproduce the inverted-U relationship between cholinergic neuromodulation and integration, reported experimentally [11, 19] and reproduced in a similar whole-brain model [10, 18]. Further, functional integration matches with an increase of SNR, a common effect attributed to neuromodulatory systems [11, 22]. The reduction of the variability of the signals (higher regularity) found in our model has also been reported experimentally in resting-state fMRI, related to an increase in functional connectivity [53]; moreover the SNR, signal variability, and functional connectivity have been shown to be influenced by neuromodulatory systems [54]. Then, the reduction in oscillatory frequency observed in the model –which falls within the Theta range of EEG spectrum– has also been perceived in several cognitive tasks [55, 56], and is a consequence of the time delays as suggested by other computational studies [46, 47]. Finally, the model is capable to reproduce the previous results of Shine *et al.* [18]: the increase in phase synchronization and functional integration with neuromodulation, the reduction of the time-resolved topological variability with integration (in our case captured by the variance of the FCD), and the inverted-U relationship of the excitatory gain *α* with *E^w^* and mean *PC^w^*.

Our main result is that the action of the cholinergic system on both, the excitation of pyramidal neurons and the intra-columnar inhibitory feedback, is needed to shift from a regime of unsynchronized activity, towards a regime of coherent activity (integrated). The inclusion of the additional inhibitory-to-excitatory interneuron connection, and its modulation through the parameter *β*, facilitates the emergence of coordinated regimens of activity (compare the cases for *β* = 0 and *β* = 0.4 of Fig 4). In our model, the inhibitory gain *β* has a double effect on the dynamic of the network. On one hand, it increases the relative magnitude of the inter-columnar afferences (compared to the internal excitatory loop), and on the other hand, it provides a simple dynamical mechanism to homeostatically preserve the excitation/inhibition (E/I) balance at the node level. In fact, this balance may be considered a determinant element in the interplay between integration and segregation [31]. Interestingly, this mechanism maximizes the functional integration in a better way than a direct decrease of the internal excitatory feedback, as we evidenced with the reduction of *C*_1_ (see Fig 2 and S2 Fig for a comparison). Our model suggests that the additional inhibitory loop, modulated by the inhibitory gain *β*, constitutes an optimal solution to the dynamical control of cortical column activity.

The inhibition-mediated control of the E/I balance has been implemented in others networks models of brain activity [57]. At the whole-brain level, Deco *et al.* [58] employed a feedback inhibitory control (FIC) to preserve the E/I balance in a mean-field network connected with a human connectome, producing the best fit between the simulated and empirical sFCs matrices. Also, FIC improved the information capacity of the global network and enhanced its dynamical repertoire constituting a biophysical plausible mechanism to reproduce the macro-scale features of human brain dynamics [58]. In our model, we implemented instead a disynaptic inhibitory control [36, 37] that, in spite of leading to similar results, can present additional advantages. The inhibition included in the model (modulated by the inhibitory gain β) influences the output of pyramidal neurons when excitability is high [36]. Because the two populations of interneurons receive excitatory inputs from pyramidal cells, an increase in pyramidal excitability triggers both the feedback excitation loop and its dampening by the inhibitory interneurons. Conversely, when the pyramidal excitability decreases, the effect of the inhibitory loop between interneurons is low, and the excitatory loop can rise the excitability of pyramidal cells. This constitutes and effective mechanism to maintain an optimal E/I balance. In contrast, a direct reduction of the excitatory feedback (e.g., decreasing *C*_1_, S2 Fig), produces no compensation when pyramidal excitability is low.

Another contribution of our work is that we propose an explicit mechanism of interaction between meso-scale properties of brain activity, such as the local inhibition, with the global increase in functional integration. While the cholinergic system increases the response of pyramidal neurons to external afferences (e.g., inputs from other brain areas) through nicotinic and muscarinic receptors [21], the increase in intra-cortical inhibition could be mediated mainly by nicotinic receptors [20, 23]. At the meso-scale level, nicotinic receptors can increment interneurons activity in a similar way than the inhibitory gain *β* [20, 22, 23]. However, there is little knowledge about the specific effects of the cholinergic system at the whole-brain level. In that line, an increase in the local and global efficiency in resting-state fMRI has been reported after the administration of nicotine [59], and some nicotinic agonists have pro-cognitive effects as well in health and disease [60]. Considering the relationship between function integration and cognition [2, 3, 10], our model suggests that the possible pro-cognitive effects associated with the cholinergic system are related to a selective increase in the excitability of excitatory and inhibitory neural populations within brain areas. Thus, our computational approach –in the same spirit as Wylie *et al.* [59]– links the meso-scale consequences of inhibitory interneurons neuromodulation with the functional network topology features, at the whole-brain level.

The inverted-U relationship between neuromodulation and integration that we found in our model is a consequence of the dynamics of individual cortical columns. At the node level, the parameters we used in the Jansen & Rit model put the model near a Hopf supercritical bifurcation [61]. When *α* and *r*_0_ are low, the node dynamics is defined by a stable focus (a fixed point with non-monotonic convergence), and thus pyramidal outputs consist of low amplitude noisy signals. Increasing both parameters causes the bifurcation into an unstable focus within a limit cycle, with high amplitude oscillations. Increasing *α* further produces a new bifurcation (a stable focus) and the limit cycle disappear. The inhibitory gain *β* constitutes a mechanism to keep the model working between the two bifurcations points; it preserves the E/I balance as an inhibitory control loop. At first, and likewise Shine *et al.* [18], we did not find an inverted-U relationship between the filter gain –modulated by the noradrenergic system– and integration. In the brain, the analogous of our parameters *α* and *β* are modulated –in parallel– by the cholinergic system, and noradrenaline has an additional effect in increasing excitability [54], thus it is possible that the inverted-U relationship observed between neuromodulation and integration is a consequence of a complex interaction between neuromodulatory systems [18]. We confirmed this changing, in parallel, both *α* and *β* parameters (see S4 Fig). The simultaneously modulation reproduced, at least partially, the inverted-U relationship between the noradrenergic system –in our model represented by the filter gain *r*_0_– and functional integration measured as the global efficiency *E^w^*. However, as the cholinergic effects on excitatory and inhibitory systems are mediated by different receptors with different expression patterns, there can be many ways by which the integration/segregation balance is controlled by neuromodulatory systems.

Near the critical boundary, in the (*α, r*_0_) parameter space, we observe an increase in the metastability and multistability, a decrease in the FCD speed, and the FCD analysis suggests an increment in dynamical richness, a consequence of the fact that each node is poised close to a Hopf supercritical bifurcation [5]. It has been proposed that at rest brain activity operates near a bifurcation point, where segregated (uncoordinated) and integrated (coordinated) regimes alternate in time [5,8,9]. In our model, the neuromodulation facilitates the transitions from “resting-state” conditions, near the critical points, to more integrated regimes. Furthermore, integrated regimes become more stable in time, as can be noted with the decrease in *d_typ_.* Analogously, time-resolved functional connectivity analysis, in task-related fMRI recordings, indicates a decrease in the topological variability in subjects performing an N-back task; the extent of integration acts as a predictor of individual performance and response times [3]. Our model reproduces the variability observed in ”resting-state” conditions, near the critical points in the parameter space, and the switching to integrated regimes with neuromodulation. Moreover, the effect of the inhibitory gain in preserving the E/I balance can play an important role in sustaining the metastability and multistability at rest. Indeed, the E/I balance is a key element to support criticality [62] and to modulate the integration and segregation features of the brain [45].

There is a lot of room for further progress starting from this work. Future research may consider the addition of neuromodulatory maps [38, 63] in order to take into account the heterogeneous expression of the receptors, or explore models that can reproduce the effect of other neuromodulatory systems [64] and their dynamics [65]. Other interneurons subtypes and their modulation could be included –such as fast-spiking inhibitory interneurons– to account for the faster EEG features of brain activity [66]. Additionally, the graph theoretical analysis used here only consider pairwise interactions, neglecting high-order effects that may contain important information about high dimensional functional brain interactions. Information-theoretical [67, 68] and algebraic topological approaches [57, 69, 70] may provide complementary insights of high-order interdependencies in the brain.

In summary, our results extend the neuromodulatory framework proposed by Shine [10], with the inclusion of the cholinergic neuromodulation of inhibitory interneurons. Importantly, we describe the homeostatic effect of this local inhibitory feedback loop, a meso-scale mechanism that interacts with macro-scale behavior and enables functional integration. Our findings shed light on a better understanding of neurophysiological mechanisms involved in the functional integration and segregation of the human brain activity. This line of research may have plentiful of scientific and clinical implications, as a vast body of evidence suggest that functional integration and segregation may be altered in neuropsychiatric disorders [31,71–73]. Therefore, understanding the neuromodulatory mechanisms that underlie the imbalances of integration and segregation may lead to future progress in pharmacological treatments.

## Materials and methods

### Whole-Brain Neural Mass Model

To simulate neuronal activity we used a reparameterized version of the Jansen & Rit neural mass model [32]. In this model, a cortical column consists of a population of pyramidal neurons with projections to other two populations: excitatory and inhibitory interneurons, which project back to the pyramidal population. Additionally, the pyramidal neurons receive an external stimulus *p*(*t*), whose values are taken from a Gaussian distribution with mean *μ* = 2 Hz and standard deviation *σ* = 2.25. Different values of *σ* were explored and our results are mostly consistent for 1.5 < *σ* < 5.

The dynamical evolution of the three populations within the cortical column is modeled by two blocks each. The first transforms the average pulse density in average postsynaptic membrane potential (which can be either excitatory or inhibitory) (Fig 1A). This block, denominated post synaptic potential (PSP) block, is represented by an impulse response function

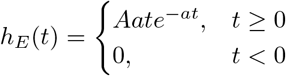

for the excitatory outputs, and

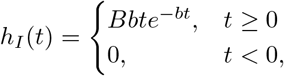

for the inhibitory ones. The constants *A* and *B* define the maximum amplitude of the PSPs for the excitatory (EPSPs) and inhibitory (IPSPs) cases respectively, while *a* and *b* represent the inverse time constants for the excitatory and inhibitory postsynaptic action potentials, respectively. The second block transforms the postsynaptic membrane potential in average pulse density, and is given by a sigmoid function of the form

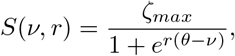

with *ζ_max_* as the maximum firing rate of the neuronal population, *r* the slope of the sigmoid function, and *θ* the half maximal response of the population.

To study the effect of the neuromodulatory systems at the macro-scale level, we included long-range pyramidal-to-pyramidal neurons and short-range inhibitory to excitatory interneurons couplings, to mimic the effects of neuromodulation through the excitatory and inhibitory gain parameters, respectively. This short-range coupling between interneurons, well described at the meso-scale level [36, 37], constitutes a modification of the original equations. In the model presented in the Fig 1A, each node *i* ∈ *N*, with *N* = [1... *n*] as the set of all nodes of the network, represents a single brain area. The nodes are connected by a structural connectivity matrix *M* (Fig 1B). This matrix is derived from a human connectome [38] parcellated in *n* = 90 cortical and subcortical regions with the automated anatomical labelling (AAL) atlas [39]; the matrix is undirected and takes values between 0 and 1. Because long-range connections are mainly excitatory [40,41], only links between the pyramidal neurons of a node i with pyramidal neurons of a node j are considered. The model includes a speed constants matrix D for the inter-columnar coupling (Fig 1B). For building the matrix D, the distance between regions’ centroids defined by the AAL atlas was used. The entries of D decrease exponentially with the distance.

The overall set of equations, for a node *i*, includes the within and between nodes

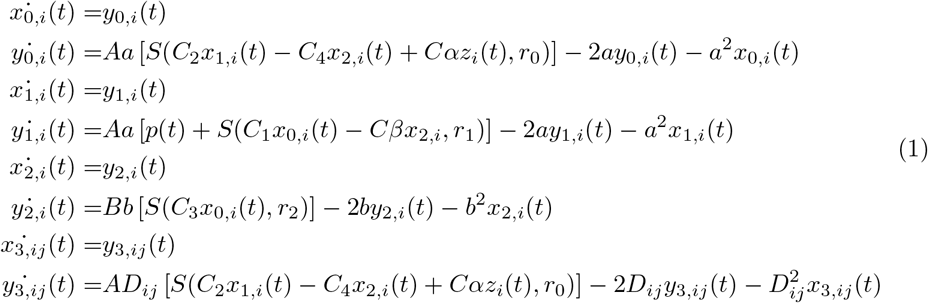

where *x*_0_, *x*_1_, *x*_2_ correspond to the outputs of the PSP blocks of the pyramidal neurons, and excitatory and inhibitory interneurons, respectively, and *x*_3,*ij*_ the output from the pyramidal neuron *j* to the column *i.* Constants *C*_1_, *C*_2_, *C*_3_ and *C*_4_ scale the connectivity between the neural populations (see Fig 1A). We used the original values of Jansen & Rit [32]: *ζ_max_* = 5 Hz, *θ* = 6 mV, *r* = 0.56 mV^−1^, *a* = 100 s^−1^, *b* = 50 s^−1^, *A* = 3.25 mV, *B* = 22 mV, *C*_1_ = *C*, *C*_2_ = 0.8C, *C*_3_ = 0.25C, *C*_4_ = 0.25C, and *C* = 135.

The overall input to the column *i* from other cortical columns is given by

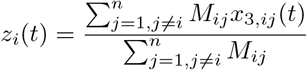

The neuronal activity of the column i is the average PSP of pyramidal neurons and characterizes the EEG-like signal in the source space; it is computed as [32]

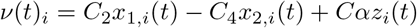

The firing rates of pyramidal neurons *ζ_i_*(*t*) = *S*(*ν*(*t*)_*i*_, *r*_0_) were used to simulate the fMRI-BOLD recordings. The parameters α, *β* and *r*_0_ account for the influence of the neuromodulatory systems (Fig 1C), as described in next subsection.

### Neuromodulation

The effects of the cholinergic system were modeled by the parameters *α* and β. The parameter *α* increases the long-rage pyramidal to pyramidal neuron coupling through the M matrix and amplifies the firing rates of the target populations [10,11]. The parameter *β* scales the short-range inhibitory to excitatory interneurons coupling, decreasing the recurrent excitation to pyramidal neurons [22]. We refer to *α* as the excitatory gain, and *β* as the inhibitory gain. In comparison with the current framework proposed by Shine [10], the novelty of our neuromodulatory approach is the inclusion of the inhibitory gain to the model. The effect of the noradrenergic system, designated as filter gain, was simulated controlling the parameter *r*_0_, which represents the sigmoid function slope of the pyramidal population. This last neuromodulatory system increases the signal-to-noise ratio and inter-columnar connectivity, promoting integration [11, 17].

### Simulation

Following Birn *et al.* [74], we ran simulations to generate the equivalent of 11 min real-time recordings, discarding the first 60 s. The system of differential equations (1) was solved with the Euler-Maruyama method, using an integration step of 1 ms. We used six random seeds which controlled the initial conditions and the stochasticity of the simulations. We simulated neuronal activity sweeping the parameters *α* ∈ [0,1], *β* ∈ [0,0.5] and *r*_0_ ∈ [0,1]. All the simulations were implemented in Python and the codes are freely available at https://github.com/vandal-uv/Neuromod2020.

### Simulated fMRI-BOLD Signals

We used the firing rates *ζ_i_*(*t*) to simulate BOLD-like signals from a generalized hemodynamic model [42]. An increment in the firing rate *ζ_i_*(*t*) triggers a vasodilatory response s¿, producing blood inflow *f_i_*, changes in the blood volume *v_i_* and deoxyhemoglobin content *q_i_*. The corresponding system of differential equations is

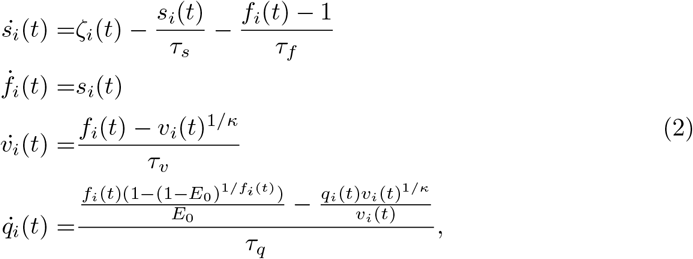

where *τ_s_, τ_f_, τ_v_* y *τ_q_* represent the time constants for the signal decay, blood inflow, blood volume and deoxyhemoglobin content, respectively. The stiffness constant (resistance of the veins to blood flow) is given by *κ*, and the resting-state oxygen extraction rate by *E*_0_. Finally, the BOLD-like signal of node *i*, denoted *B_i_*(*t*), is a non-linear function of *q_i_*(*t*) and *v_i_*(*t*)

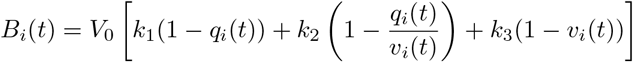

where *V*_0_ represent the fraction of venous blood (deoxygenated) in resting-state, and *k*_1_, *k*_2_, *k*_3_ are kinetic constants. We used the same parameters as in Stephan *et al.* [42]: *τ_s_* = 0.65, *τ_f_* = 0.41, *τ_v_* = 0.98, *τ_q_* = 0.98, *κ* = 0.32, *E*_0_ = 0.4, *k*_1_ = 2.77, *k*_2_ = 0.2, *k*_3_ = 0.5.

The system of differential equations (2) was solved with the Euler method, using an integration step of 1 ms. The signals were band-pass filtered between 0.01 and 0.1 Hz with a 3rd order Bessel filter. These BOLD-like signals were used to build the functional connectivity (FC) matrices from which the subsequent analysis of functional network properties is performed using tools from graph theory.

### Global Phase Synchronization

As a measure of global synchronization we calculated the Kuramoto order parameter *R*(*t*) [43] of the EEG-like signals *ν*(*t*) derived from the Jansen & Rit model. The raw signals were filtered with a 3rd order Bessel band-pass filter using their frequency of maximum power (usually between 4 and 10 Hz) ±3 Hz. Then, the instantaneous phase *ϕ*(*t*) was obtained with the Hilbert transform.

The global phase synchrony is computed as:

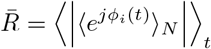

where *ϕ_i_*(*t*) is the phase of the oscillator *i* over time, 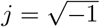 the imaginary unit, |•| denotes the module, 〈〉_*N*_ denotes the average over all nodes, and 〈〉_*t*_ the average over time. A value of 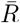 equal to 1 indicates perfect in-phase synchronization of all the set *N* of oscillators, while a value equal to 0 indicates total asynchrony.

Metastability *χ_R_*, a measure of dynamical richness, is the variance of *R*(*t*) [50]

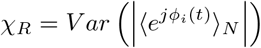

### Signal-to-noise ratio

The oscillation frequency *ω_i_* of each node *i* was computed as the peak frequency of its power spectral density function denoted *PSD*(*ω*). This function was calculated using the Welch’s method [75], with 20 s time windows overlapped by 50%.

We calculated the average signal-to-noise ratio (SNR) overall raw signals, using the *PSD*(*ω*) function of each node and excluding the 2nd to 5th harmonics [76]. For a node *i*, the signal power, *P_signal_*, was measured as the area under the curve of *PSD*(*ω*) within *ω_i_* ± 1H*z*. Noise power, *P_noise_*, corresponds to the area under the curve of *PSD*(*ω*) outside the ±1H*z* window. Then the SNR was calculated as

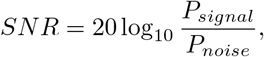

The SNR was computed for each node *i* and we reported the average overall nodes.

### Regularity

We computed the regularity index in 20 s epochs of the raw EEG-like signals *ν*(*t*) as described by Malagarriga *et al.* [48]:

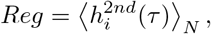

with 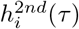 being the second peak of the autocorrelation function for the node *i* and 〈•〉_*N*_ the average over nodes. If the time series is purely periodic and has no noise, *h^2nd^*(*τ*) is equal to 1 (the signal is periodic and regular). Chaotic and noisy time series shift *h^2nd^*(*τ*) to lower values.

### Functional Connectivity and Graph Thresholding

The static Functional Connectivity (sFC) matrices were built from pairwise Pearson’s correlations of the entire BOLD-like time series. Instead of employing an absolute or proportional thresholding, we thresholded the sFCs matrices using Fourier transform (FT) surrogate data [77] to avoid the problem of introducing spurious correlations [78]. The FT algorithm uses a phase randomization process to destroy pairwise correlations, preserving the spectral properties of the signals (the surrogates have the same power spectrum as the original data). We generated 500 surrogates time series of the original set of BOLD-like signals, and then built the surrogates sFCs matrices. For each one of the (*n*^2^ – *n*)/2 possible connectivity pairs (with *n* = 90) we fitted a normal distribution of the surrogate values. Using these distributions we tested the hypothesis that a pairwise correlation is higher than chance (that is, the value is at the right of the surrogate distribution). To reject the null hypothesis, we selected a *p*-value equal to 0.05, and corrected for multiple comparisons with the FDR Benjamini-Hochberg procedure [79] to decrease the probability of make type I errors (false positives). The entries of the sFC matrix associated with a *p*-value less than 0.05 were set to 0. The result is a thresholded, undirected, and weighted (with only positive values) sFC matrix.

### Integration and Segregation

Integration and segregation were evaluated over the thresholded sFC matrices. We employed the weighted versions of transitivity [80] and global efficiency [81] to measure integration and segregation, respectively. A detailed description of the metrics used can be found in Rubinov & Sporns [44]. The transitivity (similar to the average clustering coefficient) counts the fraction of triangular motifs surrounding the nodes (the equivalent of counting how many neighbors are also neighbors of each other), with the difference that it is normalized collectively. It is defined as

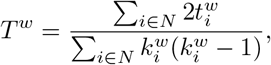

being *N* the set of all nodes of the network with n number of nodes, 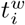 the geometric average of the triangles around the node *i*, and 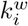 the node weighted degree. The supra-index *w* is used to refer to the weighted versions of the topological network measures. On the other hand, the global efficiency is a measure of integration based on paths over the graph: it is defined as the inverse of the average shortest path length. This metric is computed as

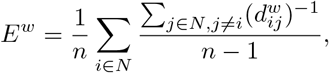

where 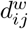 is the shortest path between the nodes *i* and *j*.

We also calculated other two measures of integration and segregation: the participation coefficient *PC^w^* and modularity *Q^w^*, respectively, both based on the detection of the network’s communities [44]. The detection of so-called communities or network modules in the thresholded sFC matrix, was based on the Louvain’s algorithm [82, 83]. The algorithm assigns a module to each node in a way that maximizes the modularity (3). We used the weighted version of the modularity [84] defined as

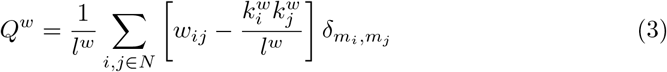

where *w_ij_* is the weight of the link between *i* and *j*, *l^w^* is the total number of weighted links of the network, *m_i_* (*m_j_*) the module of the node *i* (*j*). The Kronecker delta *δ_mi,mj_* is equal to 1 when *m_i_* = *m_j_* (that is, when two nodes belongs to the same module), and 0 otherwise. Because the Louvain’s algorithm is stochastic, we employed the consensus clustering algorithm [85]. We ran the Louvain’s algorithm 200 times with the resolution parameter set to 1.0 (this parameter controls the size of the detected modules; larger values of this parameter allows the detection of smaller modules). Then, we built an agreement matrix *G*, in which an entry *G_ij_* indicates the proportion of partitions in which the pairs of nodes (*i, j*) share the same module (so, the entries of *G* are bounded between 0 and 1). Then, we applied an absolute threshold of 0.5 to the matrix *G*, and ran again the Louvain’s algorithm 200 times using *G* as input, producing a new consensus matrix *G*’. This last step was repeated until convergence to an unique partition.

Finally, we computed the weighted version of the participation coefficient [86]. This metric quantifies, for each individual node, the strength of between-module connections respect to the within-module connections, and is defined as

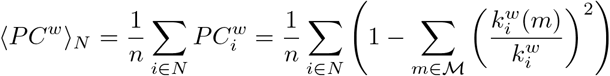

where 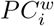 is the weighted participation coefficient for the node *i*, and 〈*PC^w^*〉_*N*_ is the average overall nodes. The functional network analysis was done in Python using the Brain Connectivity Toolbox [44].

### Functional Connectivity Dynamics

To test the hypothesis that dynamical variability peaks on the critical boundary, we performed a Functional Connectivity Dynamics (FCD) analysis over the filtered BOLD-like signals. The FCD matrix captures the evolution of FCs patterns and, consequently, the dynamical richness of the network [8, 9]. We used the sliding window approach [8,51] depicted in the Fig 6. Window length was set to 100 s with a displacement of 2 s between consecutive windows (Fig 6A). The length was chosen on the basis of the lower limit of the band-pass filter (0.01 Hz), in order to minimize spurious correlations [87]. For each window, a FC matrix was calculated from the pairwise Pearson’s correlations of BOLD-like signals (neglecting negative values), thus we obtained 251 weighted and undirected FCs matrices from the 600 s simulated BOLD-like signals (Fig 6B).

The upper triangle of each FC matrix is unfolded to make a vector, and the FCD is built by calculating the Clarkson angular distance 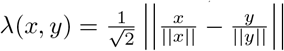 [88] between each pair of FCs (Fig 6C)

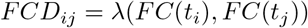

The variance of the values in the upper triangle of the FCD, with an offset of *τ* = 100 s from the diagonal (e.g., the variance of the histograms of Fig 6D), is taken as a measure of dynamical richness [51].

The speed of the FCD was measured as described by Battaglia *et al.* [52]. We computed the histogram of FCD values through a straight line from FCD(*τ*, 0) to FCD(*t_max_*, *t_max_* – *τ*), with *t_max_* = 600 s as the total time-length of the signals and *τ* = 100 s. The median of the histogram distribution corresponded to the typical FCD speed *d_typ_*. Values closer to 1 indicate a constant switching of states, and values closer to 0 correspond to stable FCs patterns.

## Supporting information

**S1 Fig. Alternative measures of network segregation and integration in the (*α,β*) parameter space. A)** Mean participation coefficient *PC^w^* (integration) and transitivity *T^w^* (segregation). **B-C)** Transitions in the direction of *α* and *β* axes. Dashed lines represent critical points.

**S2 Fig. Signal and network features in the (*α,C*_1_) parameter space. A)** Average phase synchrony 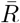, mean oscillatory frequency *ω* of EEG-like signals, global efficiency *E^w^* (integration) and modularity *Q^w^* (segregation) of the graphs derived from the sFCs of the BOLD-like signals. **B)** Transitions in the direction of *α* axis, for a fixed *C*_1_ = 0. **C)** Transitions in the direction of *C***1** axis, for a fixed *α* = 0.5. Dashed lines represent critical points.

**S3 Fig. Alternative measures of network segregation and integration in the (*α, r*_1_) parameter space. A-B)** Mean participation coefficient *PC^w^* (integration) and transitivity *T^w^* (segregation) with **A)** *β* = 0 and **B)** *β* = 0.4. **C-D)** Transitions in the direction of *α* and *r*_0_ axes. Dashed lines represent critical points.

**S4 Fig. Simultaneously effect of *α, β* and *τ*_0_ in signal and network features. A)** Average phase synchrony 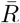, mean oscillatory frequency *ω* of EEG-like signals, global efficiency *E^w^* (integration) and modularity *Q^w^* (segregation) of the graphs derived from the sFCs of the BOLD-like signals. **B)** Transitions in the direction of *α* axis, for a fixed *r*_0_ = 0.5 mV^−1^. **C)** Transitions in the direction of *r*_0_ axis, for a fixed *α* = 0.5. Dashed lines represent critical points. Response gains change in parallel following the relationship *β* = 0.35*α*.

## Acknowledgments

We thank to Gustavo Deco, from Pompeu Fabra University (UPF, Barcelona, Spain), who kindly provided the anatomical connectivity matrix used in the model. We also want to thank to Chiayu Chiu and Andrés Chávez, from Centro Interdisciplinario de Neurociencia de Valparaíso (CINV, Valparaíso, Chile), for their feedback and suggestions about the manuscript. This work was supported by Fondecyt Grants 1181076 (to PO) and 11181072 (to RC) and the Advanced Center for Electrical and Electronic Engineering (FB0008 ANID, Chile). The Centro Interdisciplinario de Neurociencia de Valparaíso (CINV) is a Millenium Institute supported by the Millennium Scientific Initiative (ANID). CC is funded by Beca Doctorado Nacional ANID 2018-21180995.

## Notes

### Competing Interest Statement

The authors have declared no competing interest.

