## Supplementary Figures for "Cholinergic neuromodulation of inhibitory interneurons facilitates functional integration in whole-brain models"

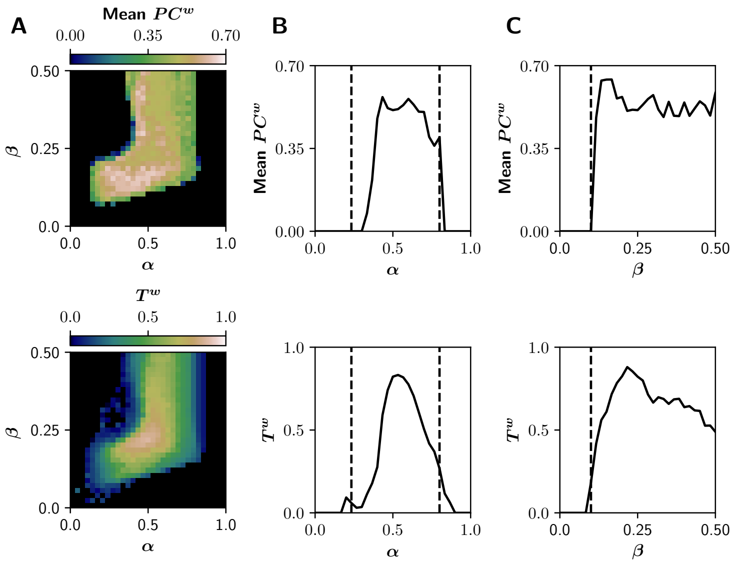


**S1 Fig. Alternative measures of network segregation and integration in the $(\alpha, \beta)$ parameter space.**
**A)** Mean participation coefficient $PC^w$ (integration) and transitivity $T^w$ (segregation). **B-C)** Transitions in the direction of $\alpha$ and $\beta$ axes. Dashed lines represent critical points.

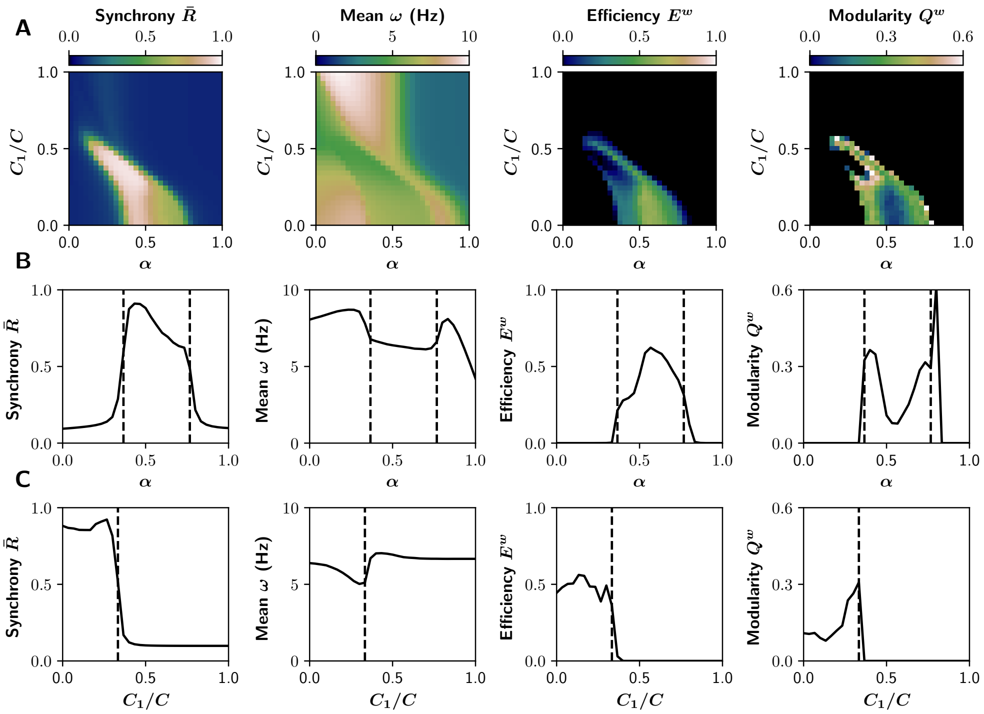


**S2 Fig. Signal and network features in the $(\alpha, C_1)$ parameter space.**
**A)** Average phase synchrony $\bar{R}$, mean oscillatory frequency $\omega$ of EEG-like signals, global efficiency $E^w$ (integration) and modularity $Q^w$ (segregation) of the graphs derived from the sFCs of the BOLD-like signals. **B)** Transitions in the direction of $\alpha$ axis, for a fixed $C_1 = 0$. **C)** Transitions in the direction of $C_1$ axis, for a fixed $\alpha = 0.5$. Dashed lines represent critical points.

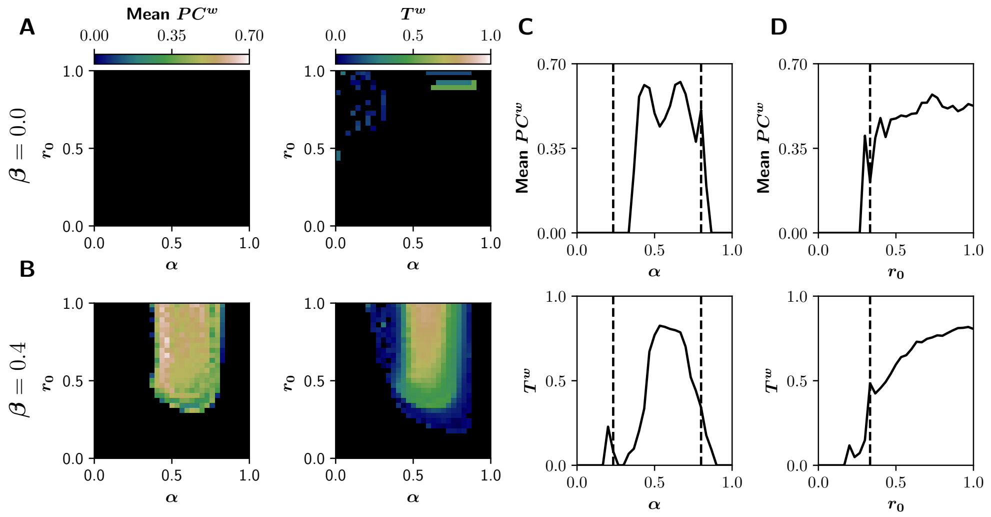


**S3 Fig. Alternative measures of network segregation and integration in the $(\alpha, r_0)$ parameter space.**

**A-B)** Mean participation coefficient $PC^w$ (integration) and transitivity $T^w$ (segregation) with **A)** $\beta = 0$ and **B)** $\beta = 0.4$. **C-D)** Transitions in the direction of $\alpha$ and $r_0$ axes. Dashed lines represent critical points.

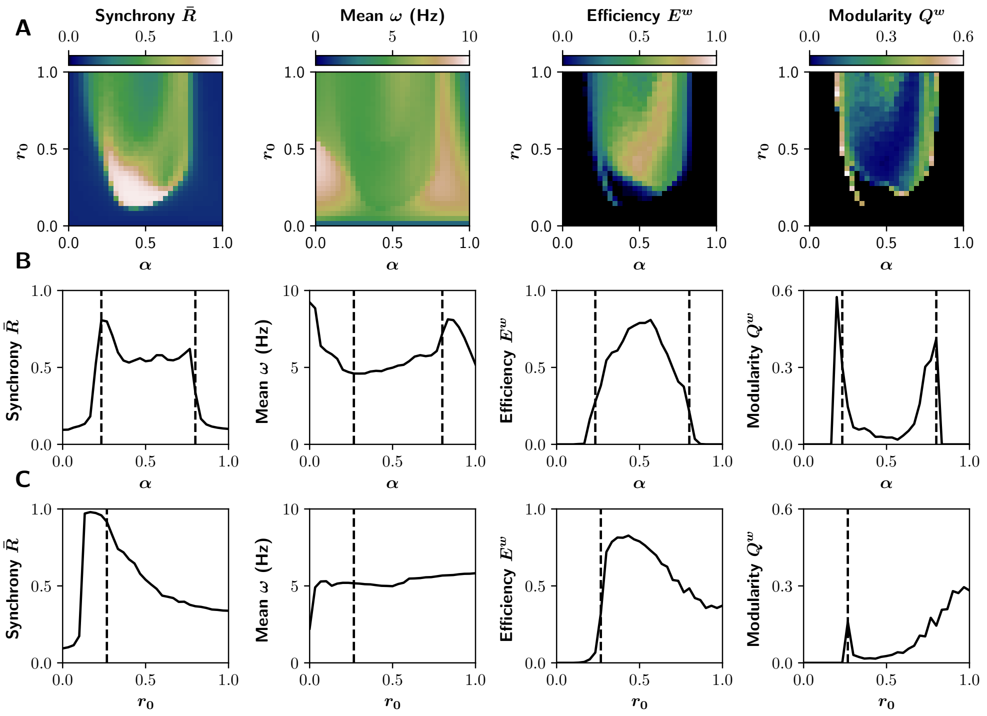


**S4 Fig. Simultaneously effect of $\alpha$, $\beta$ and $r_0$ in signal and network features.** **A)** Average phase synchrony $\bar{R}$, mean oscillatory frequency $\omega$ of EEG-like signals, global efficiency $E^w$ (integration) and modularity $Q^w$ (segregation) of the graphs derived from the sFCs of the BOLD-like signals. **B)** Transitions in the direction of $\alpha$ axis, for a fixed $r_0 = 0.5$ mV$^{-1}$. **C)** Transitions in the direction of $r_0$ axis, for a fixed $\alpha = 0.5$. Dashed lines represent critical points. Response gains change in parallel following the relationship $\beta = 0.35\alpha$.
